## Supplemental Tables and Figures for "Differential expression of RSK4 transcript isoforms in cancer and its clinical relevance"

**Figure 1**

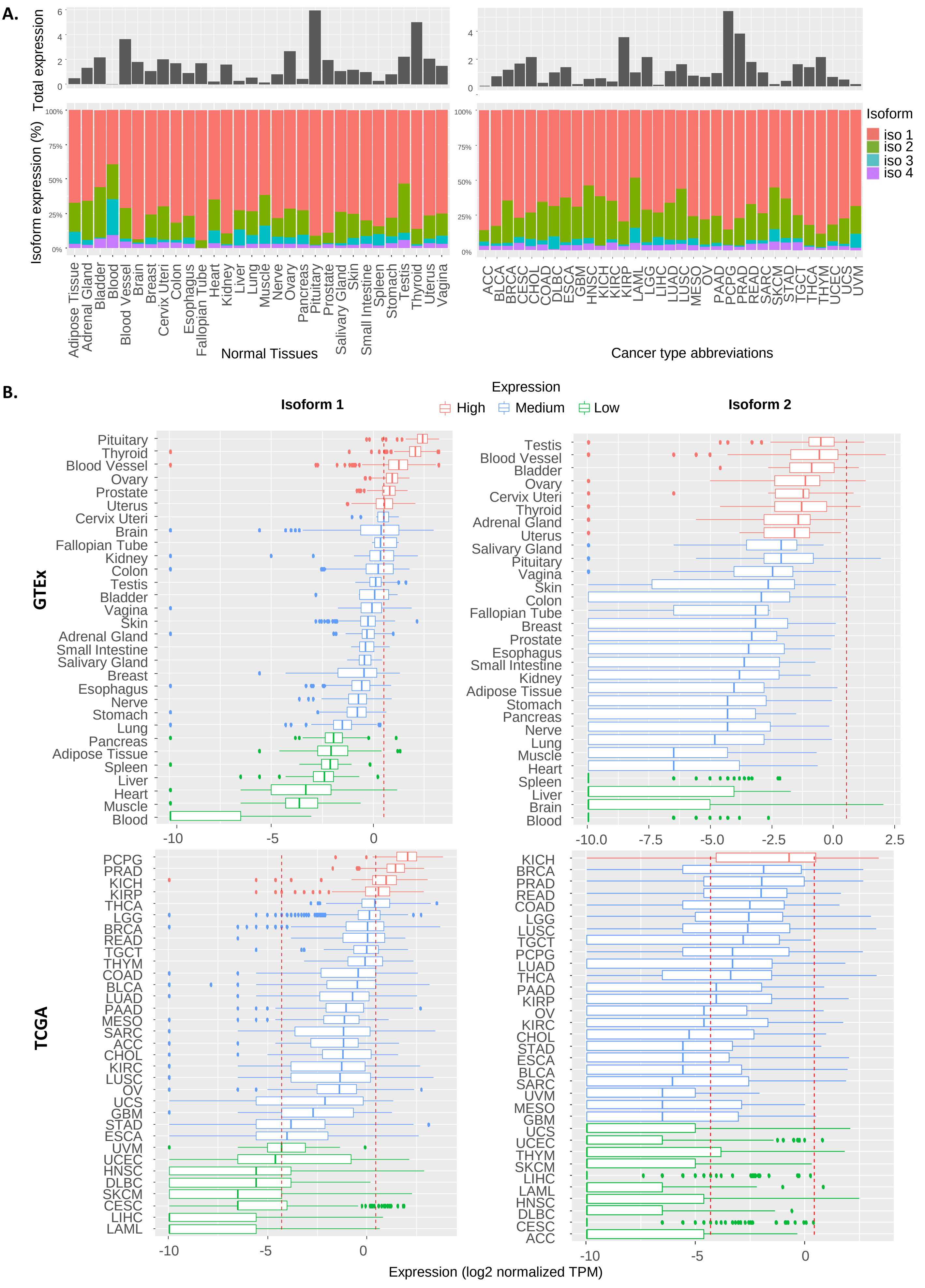

A.
Total expression
Isoform expression (%)
Normal Tissues
Cancer type abbreviations
Isoform
iso 1
iso 2
iso 3
iso 4
Adipose Tissue
Adrenal Gland
Bladder
Blood
Blood Vessel
Brain
Breast
Cervix Uteri
Colon
Esophagus
Fallopian Tube
Heart
Kidney
Liver
Lung
Muscle
Nerve
Ovary
Pancreas
Pituitary
Prostate
Salivary Gland
Skin
Small Intestine
Spleen
Stomach
Testis
Thyroid
Uterus
Vagina
ACC
BLCA
BRCA
CESC
CHOL
COAD
DLBC
ESCA
GBM
HNSC
KICH
KIRC
KIRP
LAML
LGG
LIHC
LUAD
LUSC
MESO
OV
PAAD
PCPG
PRAD
READ
SARC
SKCM
STAD
TGCT
THCA
THYM
UCEC
UCS
UVM
B.
Expression
High
Medium
Low
Isoform 1
Isoform 2
GTEx
TCGA
Expression (log2 normalized TPM)

**Figure 2**

**A.**

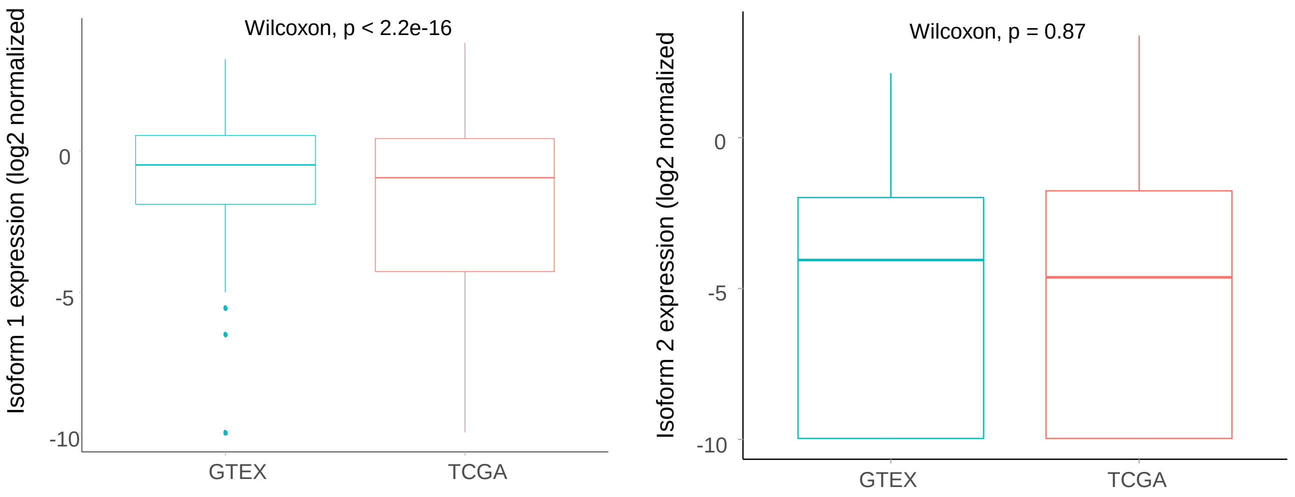


**B.**

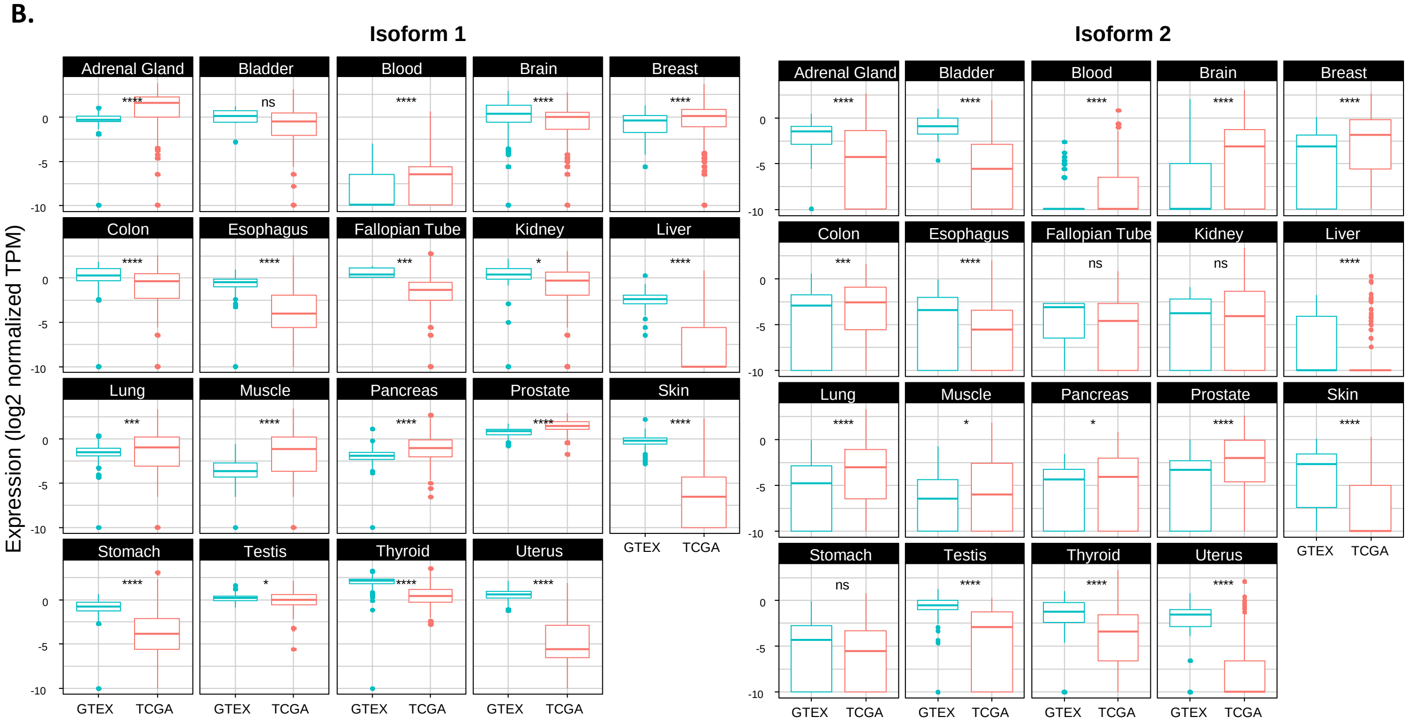


**C.**

**Isoform 1**

| GTEx | TCGA | p.adj | effsize |
|---|---|---|---|
| Skin | Skin Cutaneous Melanoma | 4.15E-156 | 2.79 |
| Uterus | Cervical & Endocervical Cancer | 1.58E-38 | 2.78 |
| Cervix Uteri | Cervical & Endocervical Cancer | 2.30E-06 | 2.71 |
| Adrenal Gland | Pheochromocytoma/Paraganglioma | 8.04E-39 | -2.50 |
| Liver | Liver Hepatocellular Carcinoma | 3.31E-37 | 2.08 |
| Thyroid | Thyroid Carcinoma | 5.52E-81 | 1.60 |
| Esophagus | Esophageal Carcinoma | 5.01E-52 | 1.50 |
| Uterus | Uterine Carcinosarcoma | 7.94E-08 | 1.48 |
| Stomach | Stomach Adenocarcinoma | 9.35E-47 | 1.48 |
| Brain | Glioblastoma Multiforme | 3.25E-54 | 1.43 |
| Cervix Uteri | Uterine Carcinosarcoma | 1.36E-02 | 1.41 |
| Blood | Diffuse Large B-Cell Lymphoma | 6.74E-13 | -1.20 |
| Prostate | Prostate Adenocarcinoma | 2.01E-20 | -1.10 |

**Isoform 2**

| GTEx | TCGA | p.adj | effsize |
|---|---|---|---|
| Cervix | Cervical & Endocervical Cancer | 1.29E-06 | 2.11 |
| Uterus | Cervical & Endocervical Cancer | 3.15E-30 | 1.88 |
| Bladder | Bladder Urothelial Carcinoma | 3.24E-05 | 1.79 |
| Cervix Uteri | Uterine Carcinosarcoma | 2.48E-04 | 1.56 |
| Adrenal Gland | Adrenocortical Cancer | 1.09E-16 | 1.44 |
| Uterus | Uterine Carcinosarcoma | 7.34E-11 | 1.36 |
| Testis | Testicular Germ Cell Tumor | 8.78E-27 | 1.34 |
| Brain | Brain Lower Grade Glioma | 1.51E-104 | -1.29 |
| Skin | Skin Cutaneous Melanoma | 7.90E-53 | 1.06 |
| Kidney | Kidney Chromophobe | 8.74E-04 | -0.93 |

**Figure 3**

**A.**

Isoform 1

| HR | LCI | UCI | PVAL | FDR | Cancer type |
|---|---|---|---|---|---|
| 0.77 | 0.71 | 0.84 | 8.18E-09 | 1.64E-08 | Brain Lower Grade Glioma |
| 0.76 | 0.63 | 0.92 | 4.46E-03 | 8.92E-03 | Kidney Chromophobe |
| 1.08 | 1.02 | 1.15 | 9.45E-03 | 1.89E-02 | Stomach Adenocarcinoma |
| 0.85 | 0.75 | 0.96 | 9.58E-03 | 1.92E-02 | Adrenocortical Cancer |
| 0.95 | 0.90 | 1.00 | 3.89E-02 | 3.89E-02 | Kidney Clear Cell Carcinoma |

Isoform 2

| HR | LCI | UCI | PVAL | FDR | Cancer type |
|---|---|---|---|---|---|
| 1.23 | 1.14 | 1.33 | 1.13E-07 | 2.26E-07 | Cervical & Endocervical Cancer |
| 0.93 | 0.90 | 0.97 | 3.41E-04 | 6.81E-04 | Kidney Clear Cell Carcinoma |
| 1.05 | 1.00 | 1.10 | 3.64E-02 | 3.64E-02 | Stomach Adenocarcinoma |
| 0.87 | 0.77 | 0.98 | 2.03E-02 | 4.05E-02 | Rectum Adenocarcinoma |

**B.**

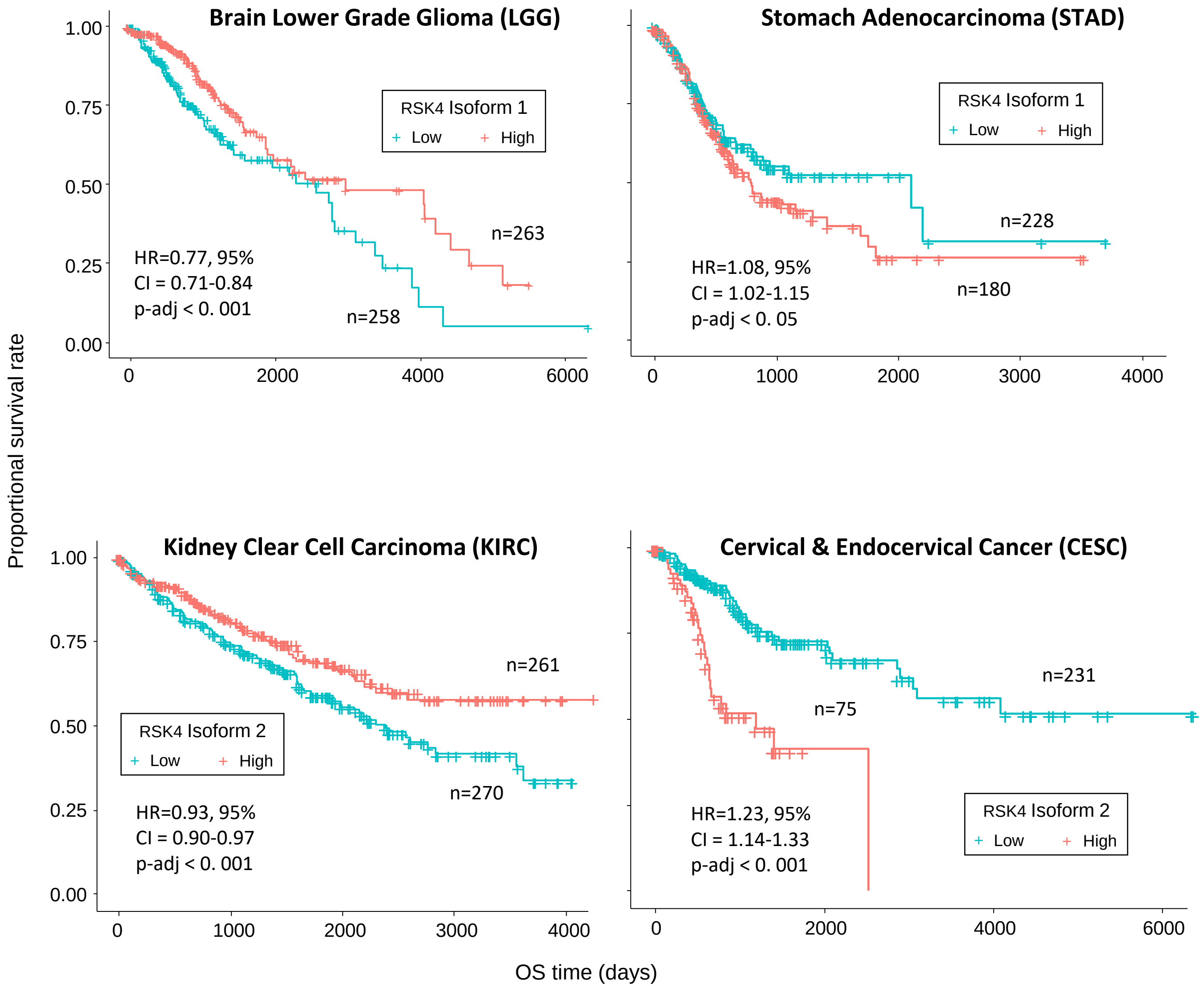

**Figure 4**

**A.**

| Cancer | Iso | Metastasis | Stage | Recurrence |
|---|---|---|---|---|
| LGG | 1 | NA | NA | -0.23075 |
| | 2 | NA | NA | |
| STAD | 1 | | | |
| | 2 | | | |
| KIRC | 1 | -0.9832 | -0.86 | -1.1652 |
| | 2 | | -1.0795 | |
| CESC | 1 | | | |
| | 2 | | 0.8746 | |

**B.**

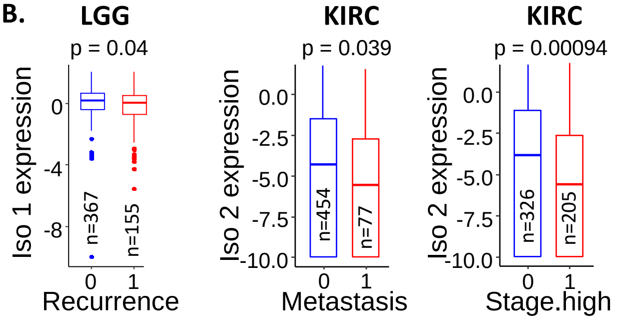


**C.**

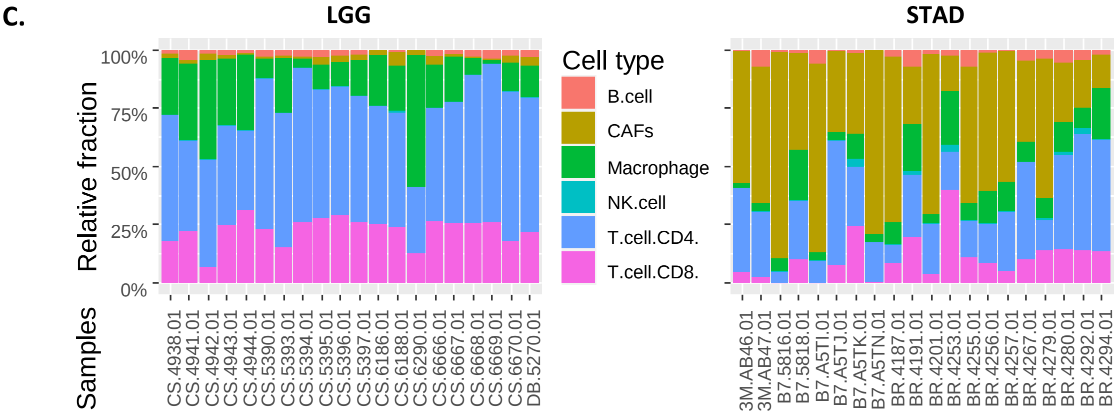


**D.**

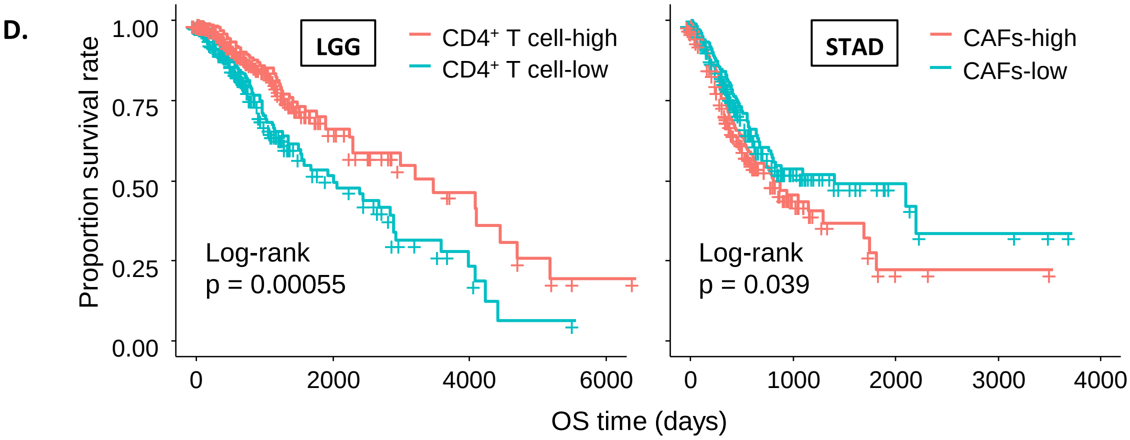


**E.**

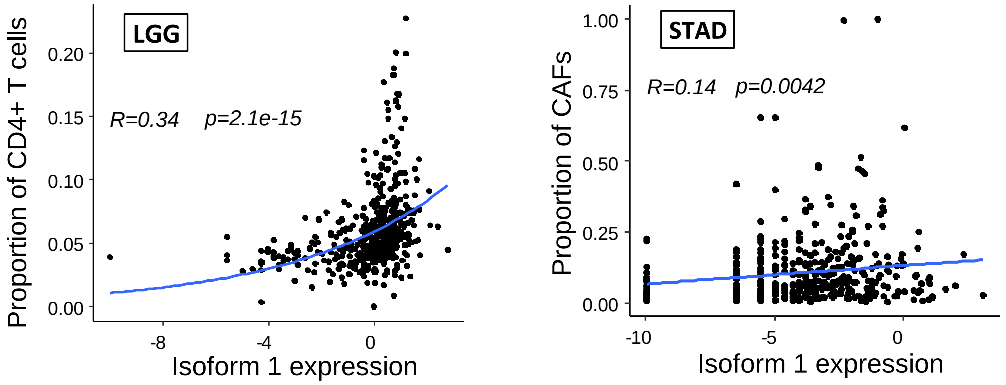


**F.**

| | Term | P-value | FDR |
|---|---|---|---|
| KEGG | TCR signaling pathway | 1.84E-04 | 0.029 |
| | Dopaminergic synapse | 5.54E-04 | 0.034 |
| | HIV1 infection | 7.26E-04 | 0.034 |
| | ErbB signaling pathway | 9.07E-04 | 0.034 |
| | Nicotine addiction | 0.001078 | 0.034 |
| Panther | Ras Pathway | 4.12E-04 | 0.009 |
| | T cell activation | 5.10E-04 | 0.009 |

**G.**

| | Term | P-value | FDR |
|---|---|---|---|
| BIOPLANET | FGFR pathway | 1.92E-04 | 0.018 |
| | Axon guidance | 2.43E-04 | 0.018 |
| | FGFR1b ligand binding and activation | 2.50E-04 | 0.018 |
| | L1-type proteins/ankyrins Interaction | 2.99E-04 | 0.018 |
| | P4-mediated oocyte maturation | 9.47E-04 | 0.046 |
| | Developmental biology | 0.0013 | 0.046 |
| | L1CAM interactions | 0.00132 | 0.046 |
| Panther | TGF-beta signaling pathway | 0.01006 | 0.235 |
| | FGF signaling pathway | 0.01382 | 0.235 |

**Supp Table 1**

| GTEX_Tissue | TCGA_Cancer types | GTEX | TCGA |
|---|---|---|---|
| Bladder | Bladder Urothelial Carcinoma | 9 | 407 |
| Breast | Breast Invasive Carcinoma | 179 | 1099 |
| Cervix & Uterus | Cervical & Endocervical Cancer<br>Uterine Carcinosarcoma | 88 | 363 |
| Brain | Glioblastoma Multiforme<br>Brain Lower Grade Glioma | 1141 | 689 |
| Fallopian Tube | Ovarian Serous Cystadenocarcinoma | 5 | 427 |
| Lung | Lung Adenocarcinoma<br>Lung Squamous Cell Carcinoma | 288 | 1013 |
| Prostate | Prostate Adenocarcinoma | 100 | 496 |
| Testis | Testicular Germ Cell Tumor | 165 | 154 |
| Esophagus | Esophageal Carcinoma | 653 | 182 |
| Pancreas | Pancreatic Adenocarcinoma | 167 | 179 |
| Kidney | Kidney Papillary Cell Carcinoma<br>Kidney Clear Cell Carcinoma<br>Kidney Chromophobe | 28 | 886 |
| Liver | Liver Hepatocellular Carcinoma | 110 | 371 |
| Muscle | Sarcoma | 1189 | 349 |
| Colon | Colon Adenocarcinoma | 308 | 290 |
| Stomach | Stomach Adenocarcinoma | 174 | 414 |
| Skin | Skin Cutaneous Melanoma | 556 | 469 |
| Thyroid | Thyroid Carcinoma | 279 | 512 |
| Blood | Acute Myeloid Leukemia<br>Diffuse Large B-Cell Lymphoma | 337 | 220 |
| Adrenal Gland | Adrenocortical Cancer<br>Pheochromocytoma & Paraganglioma | 128 | 259 |

**Supp Table 2**

### Brain Lower Grade Glioma (LGG)

| Variables | Univariate analysis | | Multivariate analysis | |
|---|---|---|---|---|
| | HR(95% CI) | adj-p-value | HR(95% CI) | adj-p-value |
| RSK4 isoform 1 | 0.77 (0.71-0.84) | 1.64e-08 *** | 0.763 (0.689-0.844) | 1.49e-07 *** |
| Family history (Yes/No) | 1.23 (0.84-1.79) | 0.283 | | |
| Gender (male vs female) | 1.12 (0.79-1.58) | 0.538 | | |
| Age (>60 vs ≤ 60) | 5.02 (3.28-7.66) | 8.3e-14 *** | 4.91 (3.19-7.58) | 5.75e-13 *** |
| Headache (Yes/No) | 0.859 (0.59-1.25) | 0.427 | | |
| Recurrence (Yes/No) | 2.71 (1.86-3.96) | 2.28e-07 *** | 2.61 (1.78-3.80) | 6.87e-07 *** |

### Stomach Adenocarcinoma (STAD)

| Variables | Univariate analysis | | Multivariate analysis | |
|---|---|---|---|---|
| | HR(95% CI) | adj-p-value | HR(95% CI) | adj-p-value |
| RSK4 isoform 1 | 1.08 (1.02-1.15) | 1.89e-02 * | 1.10 (1.03-1.18) | 0.00339 ** |
| Gender (male vs female) | 1.19 (0.849-1.67) | 0.312 | | |
| Age (>60 vs ≤ 60) | 1.66 (1.16-2.57) | 0.00582 ** | 2.12 (1.47-3.07) | 6.67e-05 *** |
| Stage.high (I&II vs III&IV) | 1.85 (1.33-2.57) | 0.000234 *** | 1.60 (1.14-2.25) | 0.00692 ** |
| Recurrence (Yes/No) | 2.50 (1.82-3.44) | 1.35e-08 *** | 2.33 (1.69-3.20) | 1.89e-07 *** |
| Metastasis (Yes/No) | 2.32 (1.36-3.96) | 0.00198 ** | 2.31 (1.32-4.02) | 0.00322 ** |

### Kidney Clear Cell Carcinoma (KIRC)

| Variables | Univariate analysis | | Multivariate analysis | |
|---|---|---|---|---|
| | HR(95% CI) | adj-p-value | HR(95% CI) | adj-p-value |
| RSK4 isoform 2 | 0.93 (0.90-0.97) | 6.81e-04 *** | 0.943 (0.906-0.981) | 0.0038 ** |
| Gender (male vs female) | 0.949 (0.696-1.29) | 0.742 | | |
| Age (>60 vs ≤ 60) | 1.83 (1.34-2.49) | 0.000128 *** | 1.67 (1.22-2.27) | 0.0012 ** |
| Stage.high (I&II vs III&IV) | 3.74 (2.73-5.13) | < 2e-16 *** | 2.59 (1.78-3.76) | 6.40e-07 *** |
| Recurrence (Yes/No) | 0.298 (0.110-0.805) | 0.017 * | 0.168 (0.0617-0.458) | 0.0005 *** |
| Metastasis (Yes/No) | 4.32 (3.16-5.90) | < 2e-16 *** | 2.62 (1.81-3.79) | 3.48e-07 *** |

### Cervical & Endocervical Cancer (CESC)

| Variables | Univariate analysis | | Multivariate analysis | |
|---|---|---|---|---|
| | HR(95% CI) | adj-p-value | HR(95% CI) | adj-p-value |
| RSK4 isoform 2 | 1.23 (1.14-1.33) | 2.26e-07 *** | 1.21 (1.12-1.32) | 5.46e-06 *** |
| Age (>60 vs ≤ 60) | 1.83 (0.545-1.10) | 0.0196 * | 1.29 (0.747-2.21) | 0.365 |
| Stage.high (I&II vs III&IV) | 2.41 (1.49-3.93) | 0.000371 *** | 1.93 (1.16-3.20) | 0.0111 * |
| Recurrence (Yes/No) | 4.70 (2.95-7.49) | 6.77e-11 *** | 4.39 (2.74-7.03) | 7.33e-10 *** |
| Metastasis (Yes/No) | 2.28 (0.83-6.29) | 0.11 | | |

**Supp Fig 1: Kruskal Wallis tests** – compare isoform 1 & 2 expression among both normal and tumour samples

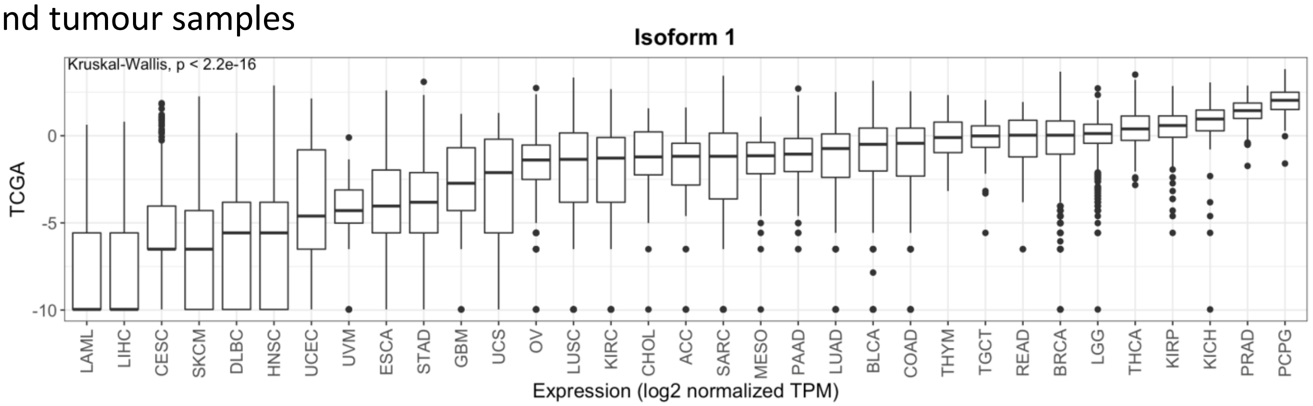


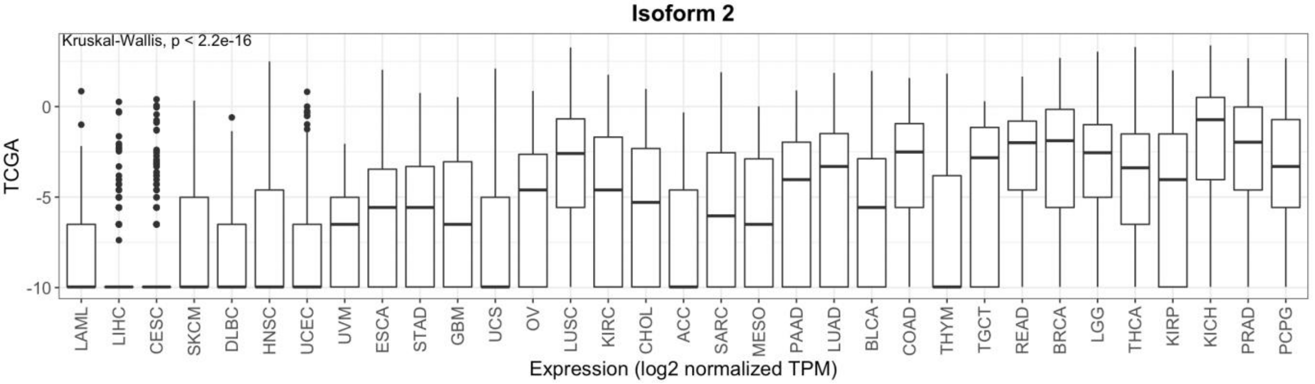


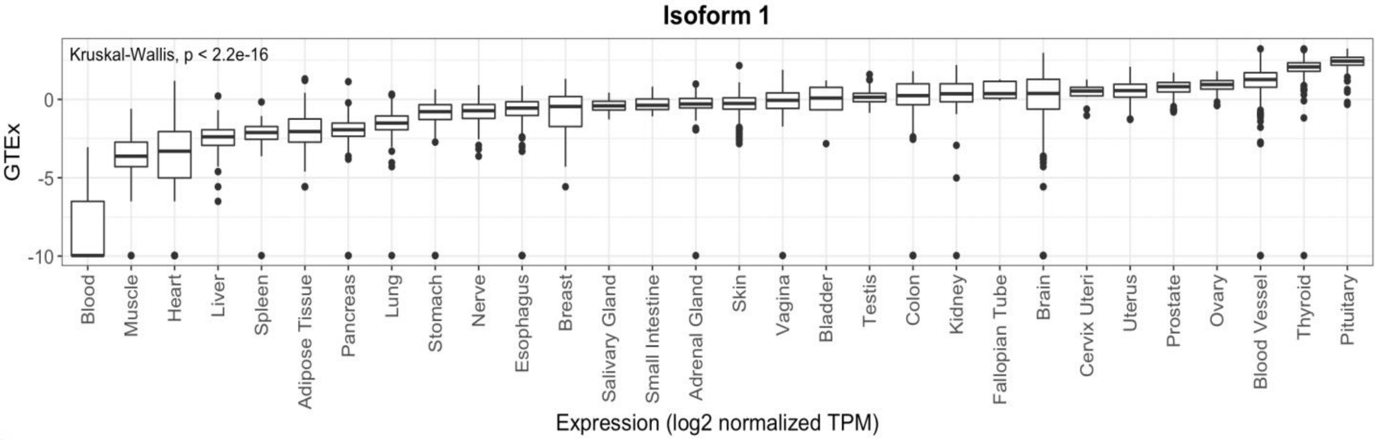


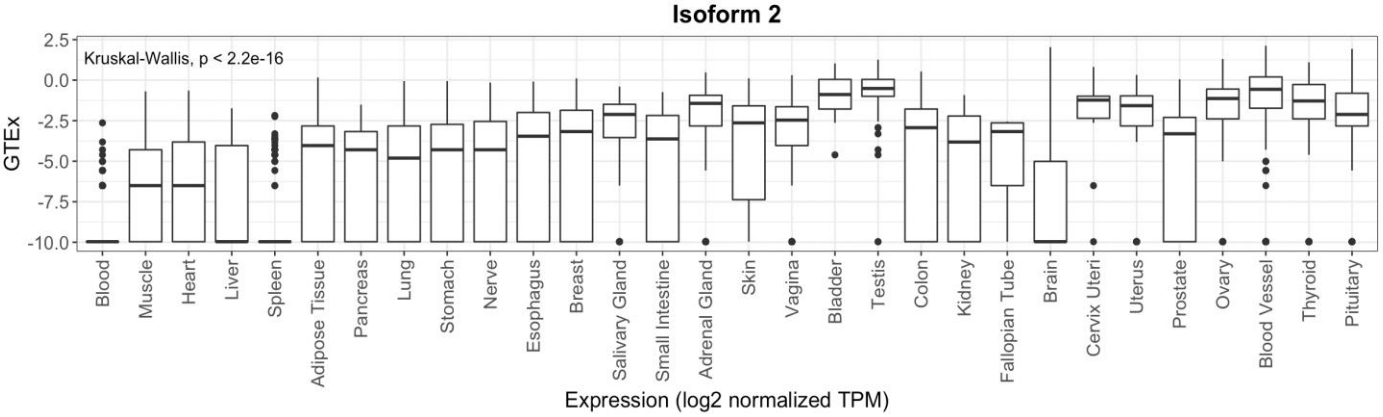

**Supp Fig 2**

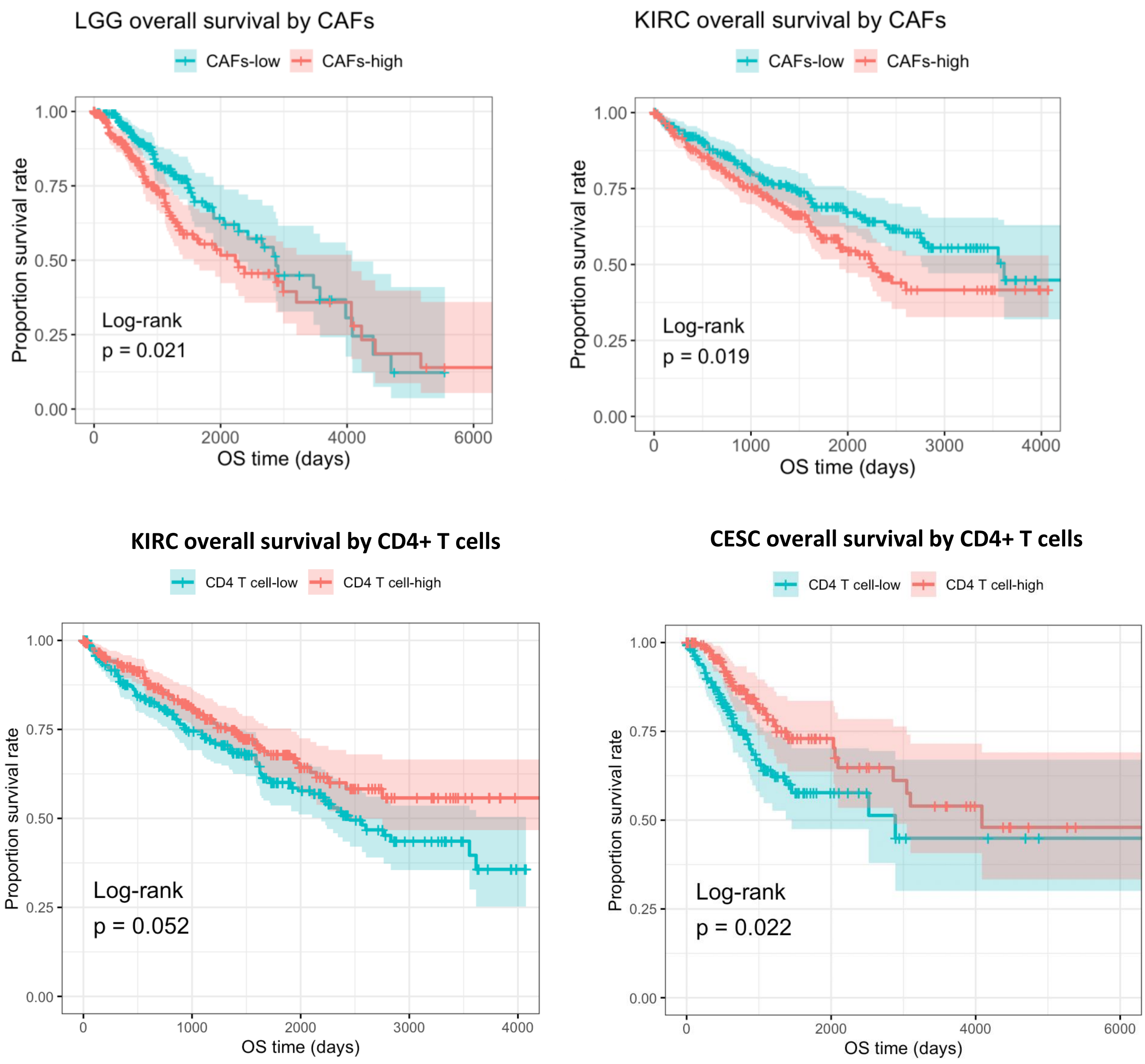
LGG overall survival by CAFs
CAFs-low
CAFs-high
Proportion survival rate
1.00
0.75
0.50
0.25
0.00
Log-rank
p = 0.021
0
2000
4000
6000
OS time (days)
KIRC overall survival by CAFs
CAFs-low
CAFs-high
Proportion survival rate
1.00
0.75
0.50
0.25
0.00
Log-rank
p = 0.019
0
1000
2000
3000
4000
OS time (days)
KIRC overall survival by CD4+ T cells
CD4 T cell-low
CD4 T cell-high
Proportion survival rate
1.00
0.75
0.50
0.25
0.00
Log-rank
p = 0.052
0
1000
2000
3000
4000
OS time (days)
CESC overall survival by CD4+ T cells
CD4 T cell-low
CD4 T cell-high
Proportion survival rate
1.00
0.75
0.50
0.25
0.00
Log-rank
p = 0.022
0
2000
4000
6000
OS time (days)
